# scDeepVariant: A population-informed deep learning framework for germline variant calling in single-cell RNA sequencing data

**DOI:** 10.64898/2025.12.31.696877

**Authors:** Ilia Buralkin, Hu Chen, Zhandong Liu, Junseok Park

## Abstract

Single-cell RNA sequencing (scRNA-seq) provides unprecedented resolution of cellular heterogeneity while also capturing information on germline genetic variation, but accurate variant calling remains limited by sparse coverage, allelic imbalance, and RNA-specific artifacts. Existing single-cell methods, including cellSNP, scAllele, and Monopogen, address some of these challenges, yet either suffer from low sensitivity and precision or rely on linkage disequilibrium (LD) priors that restrict performance on rare variants. Here, we introduce scDeepVariant (scDV), a deep learning-based framework adapted from DeepVariant and trained on paired whole-genome sequencing (WGS) and single-nucleus RNA sequencing (snRNA-seq) data. We show that scDV can be effectively trained on sparse single-cell data and that augmenting models with allele frequency information from gnomAD or the 1000 Genomes Project consistently improves performance. Across benchmarks, scDV with allele frequency channels achieved higher precision and recall than standard six-channel configurations, surpassing Monopogen at coverage depths above 10× and demonstrating a pronounced advantage in rare variant detection, where LD-based refinement is most limited. These results establish scDV as a robust alternative for germline variant discovery from scRNA-seq and highlight the broader value of integrating population-scale information into deep learning frameworks for transcriptomic variant calling.

## Introduction

Single-cell RNA sequencing (scRNA-seq) has transformed transcriptomics by enabling large-scale characterization of cellular heterogeneity across tissues ^[1]^ and disease contexts ^[2]^. While primarily designed to profile gene expression states, scRNA-seq also generates read alignments that contain information about germline genetic variants. Traditionally, germline variant discovery relies on whole-genome sequencing (WGS) or whole-exome sequencing (WES) of bulk DNA samples. Leveraging scRNA-seq for germline variant calling offers the potential to expand the utility of already-generated datasets without additional sequencing, particularly for transcribed regions captured in transcriptomic assays.

Despite these advantages, accurate variant calling from scRNA-seq remains challenging. Coverage is sparse and highly uneven because only a subset of the transcriptome is captured in each cell. Technical and biological factors compound this problem: (i) 5^*′*^- and 3^*′*^-end enrichment caused by library preparation biases; (ii) cell-type–specific expression patterns that cause a large dynamic range in read depth; (iii) allelic imbalance and allele-specific expression; (iv) errors introduced during reverse transcription, amplification, or sequencing; and (v) splicing and RNA-editing events that obscure true DNA variants. As a result, applying bulk DNA variant callers such as SAMtools ^[3]^, GATK ^[4]^, FreeBayes, or Strelka2 ^[5]^ to scRNA-seq typically recovers fewer than 8% of true variants, even in full-length SMART-seq2 data, and performs worse in droplet-based assays ^[6]^.

To overcome these limitations, methods tailored for single-cell data have been developed. cellSNP relies on direct read pileups against specified SNP sites (or genome-wide in de novo mode), using libraries such as pysam to count alleles per cell or sample ^[7]^. scAllele, in contrast, employs a local reassembly strategy: it constructs de Bruijn graphs for each region with overlapping reads, and candidate variants are scored with a generalized linear model that incorporates base quality, allelic ratio, stutter noise, and splicing context ^[8]^. While these approaches attempt to address scRNA-specific biases, benchmarking shows that both suffer from low sensitivity and precision ^[6]^. More recently, Monopogen has emerged as the state-of-the-art single-cell variant caller ^[6]^. By leveraging linkage disequilibrium (LD) from external reference panels such as the 1000 Genomes Project (1KGP) ^[9]^, Monopogen achieves high genotyping accuracy (>95% for variants present in the panel) while also detecting putative somatic variants through allele cosegregation patterns at the cell population level. However, this reliance on LD refinement inherently reduces sensitivity to rare germline variants (<0.01% allele frequency), which are often absent or underrepresented in population panels.

An alternative paradigm is offered by deep learning. DeepVariant, originally developed for WGS and WES, applies convolutional neural networks to image-like pileup representations of sequencing reads and has consistently outperformed traditional statistical and algorithmic callers such as GATK, Strelka, and DRAGEN ^[10]^. Unlike these handcrafted-feature approaches, DeepVariant learns directly from sequence features embedded in images, enabling it to capture subtle patterns that distinguish true variants from artifacts. Despite the lower and more uneven coverage of RNA-seq data, the same principles can be applied to learn RNA-specific artifacts such as splicing junctions, editing events, and allelic imbalance. Recent work demonstrated this by adapting DeepVariant to bulk RNA-seq, first using GIAB RNA with matched truth genotypes and later scaling to “silver truth” training sets derived from GTEx samples with matched WGS ^[11]^. Although the silver truth labels were less accurate than GIAB, the vastly larger sample size and tissue diversity enabled models that surpassed GIAB-trained counterparts, showing that DeepVariant can be effectively trained for transcriptomic variant calling when sufficient paired RNA–DNA data are available.

Here, we introduce scDeepVariant (scDV), a deep learning–based framework adapted from DeepVariant for germline variant discovery in scRNA-seq data. Leveraging paired WGS and single-nucleus RNA sequencing (snRNA-seq) samples, we demonstrate that scDV can be effectively trained on sparse single-cell data, despite the lower sequencing depth characteristic of these assays. Motivated by prior evidence that population priors improve variant calling through allele frequency (AF) channels in DeepVariant for WGS/WES ^[12]^ and linkage disequilibrium–based refinement in Monopogen, we further evaluated the impact of incorporating AF as an additional input channel. Across all benchmarks, scDV models augmented with AF information consistently outperformed standard six-channel configurations. While Monopogen achieved the highest performance across all sites at the ≥3× coverage threshold, scDV surpassed it when sites were restricted to ≥10× coverage and showed the most pronounced advantage in rare variant detection, a setting in which LD-based approaches are particularly constrained. Collectively, these findings establish scDV as a promising new alternative for single-cell variant calling while highlighting the broader potential of integrating population-scale information into deep learning methods for transcriptomic variant discovery.

## Materials and Methods

### Training samples

We curated a dataset of 22 human samples from the Religious Orders Study and Memory and Aging Project (ROSMAP) cohort ^[13]^, each with matched snRNA-seq and WGS data. snRNA-seq libraries were prepared using the 10x Genomics Chromium Single Cell 3’ Reagent Kits v3 TOSS and sequenced on Illumina NovaSeq 6000 (or NextSeq 500/550 for some batches). Whole-genome sequencing was carried out on the Illumina HiSeq X Ten platform at approximately 30× coverage per sample. A detailed description is available on Synapse (Project SynID: syn3219045).

#### snRNA-seq processing

We aligned raw FASTQ files to the GRCh38 reference genome using STAR (v2.7.10a) ^[14]^ with default parameters and with `–soloType Droplet` and `–soloFeatures Gene` enabled to support single-cell quantification. Coordinate-sorted BAMs were produced directly by STAR. Sample-level BAMs were renamed and indexed using SAMtools ^[3]^, followed by a secondary sorting and re-indexing step for consistency.

#### WGS VCF preprocessing

WGS variant calls were obtained from ROSMAP joint genotyping VCFs. We separated multi-sample VCFs into individual sample-level files using `bcftools view` ^[15]^, extracting and filtering chromosomes individually. These were then concatenated per sample across chromosomes using `bcftools concat`. Because the original data were aligned to GRCh37, we performed a liftover to GRCh38 using CrossMap ^[16]^ and the `hg19ToHg38.over.chain.gz chain file`. Lifted VCFs were sorted, compressed, and indexed. To finalize the preprocessing, we normalized variant representations using `bcftools norm`, filtered for non-reference genotypes (`GT !`= 0/0), and fixed contig naming inconsistencies with custom regular expressions.

#### Coverage-based filtering

We calculated per-base coverage on snRNA-seq BAMs using mosdepth (v0.3.1) ^[17]^ and retained only regions with minimum depth ≥3× to restrict training to confidently callable regions. Resulting BED files defined high-confidence transcriptomic intervals for training and inference. All downstream variant calling from snRNA-seq was limited to these regions. Each sample was then represented by its snRNA-seq BAM and corresponding WGS-derived truth variants aligned to the same GRCh38 coordinate system.

#### Study structure and chromosome holdout

We partitioned the genome by chromosome to create the training split and ensure unbiased evaluation. Chromosomes 1 through 19 were used to train the model, while chromosomes 21 and 22 were reserved for hyperparameter tuning and validation. Chromosome 20 was completely held out from training and validation and served as a true test set for model performance evaluation. This chromosome-based partitioning strategy, which avoids random sample-level splits, ensures no leakage of genomic context between training and testing sets.

#### Evaluation dataset

We curated an independent evaluation dataset consisting of seven human samples with matched scRNA-seq and WGS or WES data to benchmark against existing tools and assess the generalization of our models beyond the ROSMAP training cohort. All seven samples were processed independently from the training set and were not used in model development, tuning, or parameter selection.

Two samples were obtained from a study of esophageal squamous-cell carcinoma by Zhang et al. ^[18]^. For the scRNA-seq data, we applied the same processing pipeline used for the ROSMAP cohort, including alignment to the GRCh38 reference genome with STAR ^[14]^. Variant calls were generated from WES data using DeepVariant’s WES model with default parameters ^[10]^ and served as ground truth. For tools requiring duplicate-marked BAMs (GATK HaplotypeCaller and Strelka2), we marked duplicates using Picard MarkDuplicates ^[19]^.

Five samples were drawn from a squamous cell carcinoma cohort published by Ji et al. ^[20]^. For scRNA-seq, we used aligned BAM files provided by the authors, originally mapped to GRCh37. To enable evaluation with tools requiring GRCh38 input (scDV and Monopogen), we lifted these BAM files to GRCh38 using CrossMap ^[16]^. For tools compatible with GRCh37 input (GATK HaplotypeCaller and Strelka2), we ran the callers on the original GRCh37 BAM files and subsequently lifted the resulting variant calls to GRCh38 for comparison. For WES ground truth, we aligned reads to GRCh38 using BWA-MEM ^[21]^, marked duplicates ^[19]^, and called germline variants using GATK HaplotypeCaller ^[4]^. All performance evaluations were conducted on GRCh38 coordinates.

### Model development

We used the DeepVariant training pipeline ^[10]^ to develop models capable of predicting germline variants directly from snRNA-seq data. To accommodate the spliced structure of RNA alignments, we enabled the –use_spliced_alignments flag during both candidate generation and training. This flag allows DeepVariant to correctly interpret intron-spanning reads by handling skipped CIGAR operations typically present in transcriptomic alignments. Using this configuration, we trained a total of five DeepVariant models.

Two of these models used the canonical six-channel pileup encoding (base identity, base quality, mapping quality, strand, variant support, and reference mismatch). One of the six-channel models was warm-started from the RNA Deep-Variant model checkpoint, while the other was initialized from the Inception v3 base architecture used by DeepVariant. The remaining three models used a seven-channel architecture by incorporating external population AF as an additional input channel. Population information was integrated through an allele matching algorithm that constructs local haplotypes to precisely match variant candidates against a reference panel ^[12]^.

Given a variant candidate *v* with genomic positions *v*_start_ and *v*_end_, the algorithm queries all cohort variants *V*_*C*_ overlapping the window [*v*_start_, *v*_end_). The complete set of variants to consider is defined as:

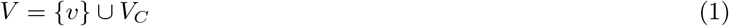

The matching window is extended to encompass all variants in *V* :

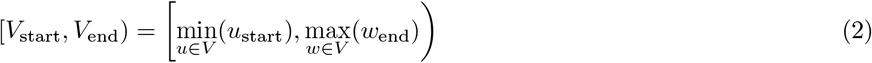

For each variant allele *a* ∈ *V*, a local haplotype *H*_*a*_ is constructed by applying the allele to the reference sequence *R* within the extended window. The AF for candidate variant *v* with alternate allele *a*_*v*_ is computed by matching haplotypes:

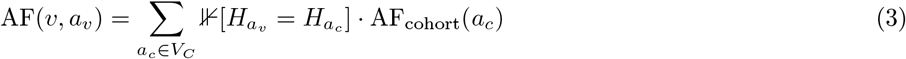

where 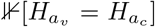 is an indicator function that equals 1 when haplotypes match perfectly, and AF_cohort_(*a*_*c*_) denotes the population AF of cohort allele *a*_*c*_ from the reference panel.

This AF information is encoded into the pileup images through a logarithmic transformation to enhance resolution for low-frequency variants. For each read overlapping the variant candidate position, the transformed frequency of its allele is used to color the corresponding pixels in the AF channel. Since individual reads may carry multiple alternate alleles with different frequencies, color intensity varies across pileup images for different variant candidates.

This approach was implemented using DeepVariant’s `–use_allele_frequency` mode, which injects cohort-level AF information into each pileup image on a per-read basis ^[12]^. All three AF models were warm-started from the Inception v3 checkpoint and differed only in the source of AF data: one model used frequencies from 1KGP ^[9]^, one used unfiltered gnomAD v3 ^[22]^, and the third used gnomAD v3 ^[22]^ filtered to include only PASS variants.

### Comparison with other SNV callers

For comparative evaluation, we ran scDV, Monopogen, GATK HaplotypeCaller ^[4]^, and Strelka2 ^[5]^ on the seven-sample evaluation dataset described above. Metrics for each sample were computed at scRNA-seq coverage of at least 3× . Monopogen was run using default germline variant calling flags. GATK HaplotypeCaller and Strelka2 were run with default settings appropriate for bulk RNA-seq data. All tools were evaluated using the same ground truth variant sets and genome-build harmonization strategy detailed in the Evaluation dataset section.

### Performance metrics

Variant calling performance was evaluated using matched WGS/WES data as ground truth. For each sample, we extracted biallelic loci with at least one alternate allele (genotypes 0/1 or 1/1) from both the called variants and ground truth sets. Variants were classified as true positives (TP) only when both genomic position and genotype matched between call sets. Variants called by the tool but absent from ground truth, or with discordant genotypes, were classified as false positives (FP). Variants present in ground truth but missed or miscalled by the tool were classified as false negatives (FN). From these counts, we computed standard classification metrics:

**Precision** measures the fraction of called variants that are correct:

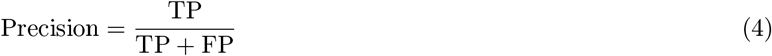

**Recall** measures the fraction of true variants that are successfully called:

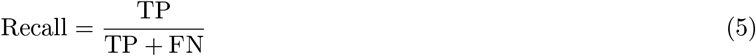

**F1-score** is the harmonic mean of precision and recall:

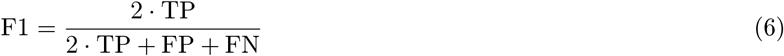

**Accuracy** measures the overall fraction of variants correctly classified:

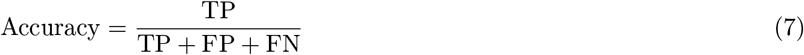

This genotype-aware evaluation framework ensures that variants are only considered correctly called when both genomic position and zygosity match the ground truth, providing a stringent assessment of variant calling performance.

### Evaluation across coverage cutoffs

We evaluated each tool across a range of minimum coverage thresholds (3×, 5×, 7×, 10×, 20×, 30×, and 50×) to assess the impact of sequencing depth on variant calling performance. For each sample, per-base coverage was calculated from the scRNA-seq BAM file using `mosdepth` ^[17]^. Both tool-called variants and ground truth variants were filtered to retain only positions meeting or exceeding each coverage threshold. Performance metrics (precision, recall, F1-score, and accuracy) were then computed for each filtered variant set as described in the Performance metrics subsection.

### Evaluation of rare variant detection

We stratified variants by their population AF using gnomAD v3 ^[22]^ to evaluate performance on rare SNVs. Only variants present in gnomAD were included in this analysis. For each AF threshold (ranging from 0.1 to 0.0001), we filtered both tool-called variants and ground truth variants to retain only variants with AF below the specified cutoff. Performance metrics (precision, recall, F1-score, and accuracy) were then computed for each AF-stratified variant set as described in the Performance metrics subsection.

## Results

### Population information improves variant calling performance in scRNA-seq data

We developed scDV by incorporating an AF channel encoded from 1KGP ^[9]^ and gnomAD v3 ^[22]^ databases, and trained it on 22 paired snRNA-seq and WGS samples from the ROSMAP cohort ^[13]^ (see Methods: Training samples). For model training, we used chromosomes 1–19 as the training set and reserved chromosomes 21–22 for hyperparameter validation, using seven-channel pileup encodings (base identity, base quality, mapping quality, strand, variant support, reference mismatch, and a channel encoding population AF) as input to an Inception-v3 CNN backbone ^[23]^ (see Methods: Model development). We trained five model variants to systematically evaluate the contribution of population information and initialization strategy, using AF channels derived from different population databases (see Methods: Model development). We then evaluated model performance across genomic regions and population AF using a chromosome 20 holdout test set from the ROSMAP dataset and observed a consistent performance gap between AF-augmented and standard models in single-cell variant calling (Figures 1B and 1C). Figure 1A shows an overview of the scDV workflow.

**Figure 1:**
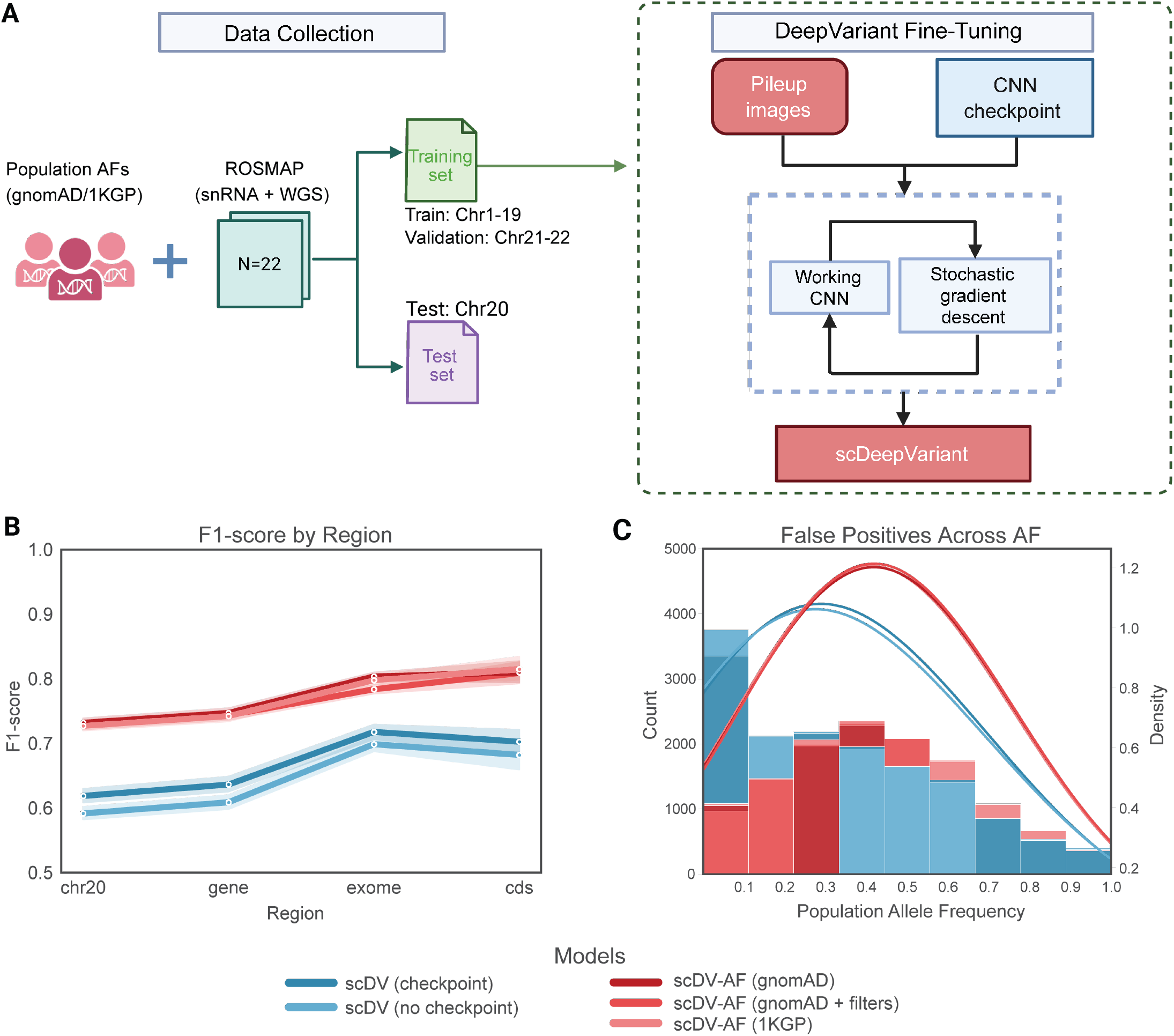
Overview of scDV training and evaluation pipeline. Figure 1A: Data collection and model fine-tuning workflow. Paired ROSMAP snRNA-seq and WGS datasets are combined with population AF from gnomAD or 1KGP, then partitioned into training (Chr1–19), validation (Chr21–22), and test (Chr20) sets. DeepVariant is fine-tuned by generating pileup images from aligned reads and training a CNN using stochastic gradient descent, initialized either from a bulk RNA-seq checkpoint or from scratch. Figure 1B: Model performance on the chromosome 20 holdout set. F1 scores across genomic regions (chr20, gene, exome, CDS) comparing AF-integrated (red) and standard six-channel (blue) models. Figure 1C: Kernel density distributions of false positive calls as a function of population AF, contrasting AF-integrated (red) and baseline (blue) models.

Figure 1B shows that AF-augmented scDV models achieved F1 scores above 0.7 across the full chromosome, gene bodies, exomes, and coding sequences, while standard six-channel models plateaued between 0.6 and 0.7. This represented absolute F1 score gains exceeding 0.10 in all tested regions, with performance on the entire chromosome 20 improving from 0.60 with six channels to 0.72 when AF was included. These improvements were robust across AF sources (1KGP, gnomAD, or gnomAD+filters), indicating that the benefit derives from population-level information rather than from specific database characteristics. The performance advantage was most pronounced for heterozygous variant calls (F1 gap >0.15 compared to <0.10 for homozygous alternate calls; Supplementary Figure 1A and 1B), where AF models achieved F1 scores of 0.65 and above versus 0.49 for standard models, reflecting how these sites are particularly vulnerable to allele dropout in single-cell data. The AF channel addresses this by providing population-level context that complements limited read evidence, an effect previously demonstrated to reduce systematic errors in low-coverage WGS variant calling ^[12]^.

Figure 1C shows the distribution of false positives by AF, illustrating how the added channel reduces systematic errors. Without AF integration, models showed false-positive rates that were at least 1.6-fold higher. AF-augmented models achieved 3-fold reductions in false positives for variants with AF < 0.1, increasing to at least 6-fold for rare variants (AF < 0.01). This effect was especially pronounced for homozygous calls, which reached more than a 51-fold reduction compared with a 3-fold reduction for heterozygous variants (Supplementary Figure 1C and 1D). The overall reduction in false positives across all AF levels demonstrates the broad utility of population context, as seen in previous WGS applications ^[12]^; the substantial improvement for rare variants, particularly for homozygous calls, indicates that these error-prone categories benefit most from AF integration.

These findings establish that population AF provides essential prior information for variant calling in low-coverage single-cell contexts, enabling scDV to resolve ambiguous calls that sequence features alone cannot confidently classify. Across genomic regions, AF integration yielded robust F1 gains exceeding 0.10, improved detection of heterozygous variants susceptible to monoallelic expression, and substantially reduced false positives at rare AF levels, supporting that population context helps to address core limitations of variant calling from sparse transcriptomic data.

### Benchmarking against state-of-the-art variant callers on independent scRNA-seq data

We benchmarked scDV on an independent cohort to demonstrate the model’s generalizability beyond the training data and to assess the effect of AF integration in single-cell variant calling. Using an external test set of seven paired scRNA-seq/WES samples held out from training, we evaluated model performance and compared it against state-of-the-art single-cell variant callers (see Methods: Evaluation dataset). Integration of population AF consistently improved both precision and recall, with AF-augmented models achieving F1 scores of 0.836–0.840. These values were within F1 of the state-of-the-art LD-based method Monopogen while substantially outperforming existing bulk RNA-seq variant-calling pipelines (Figure 2A).

**Figure 2:**
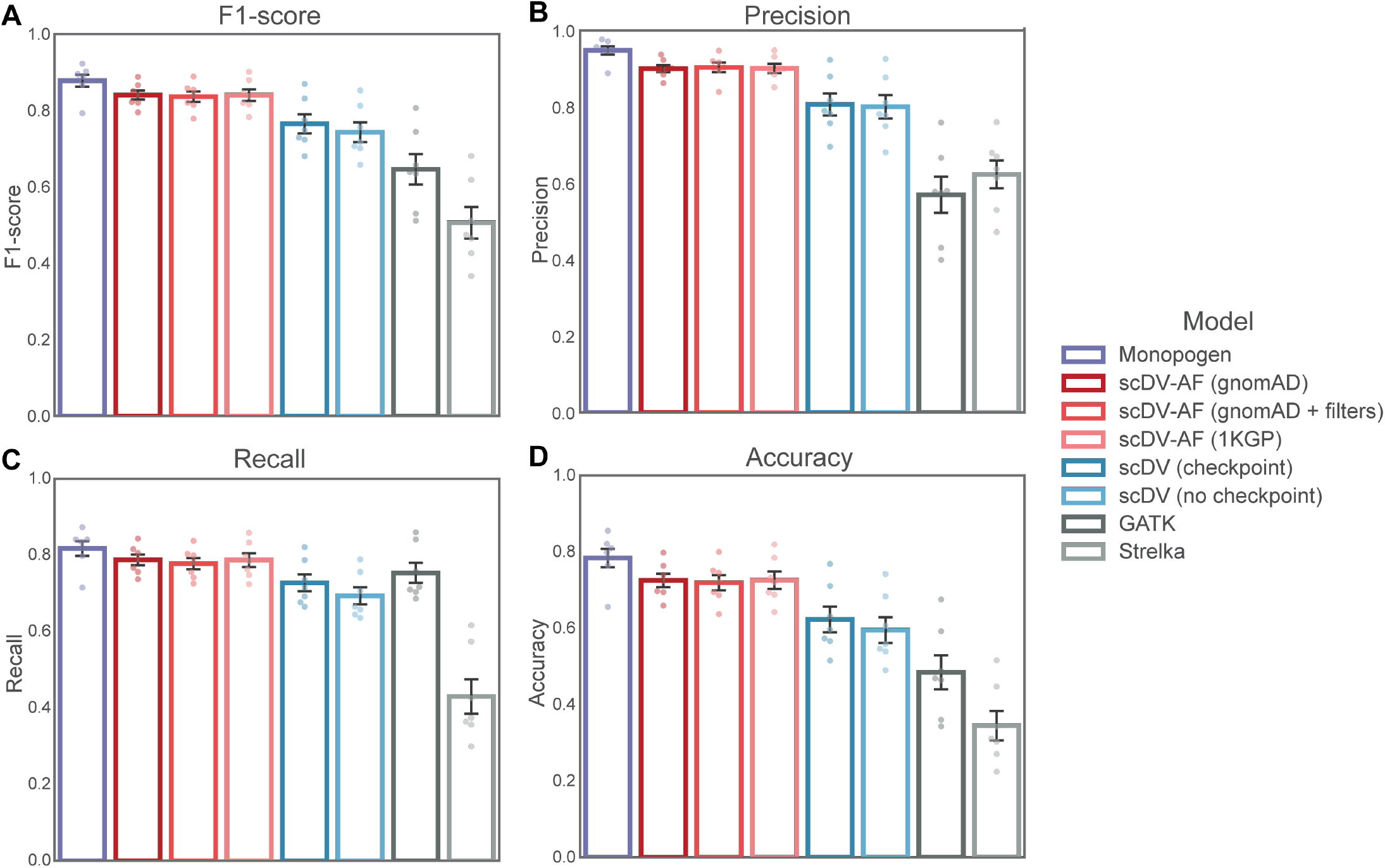
Performance comparison of 8 variant calling methods evaluated on seven independent scRNA-seq samples with matched WES ground truth. Panels show (A) precision, (B) recall, (C) F1-score, and (D) accuracy. Methods include: Monopogen (purple), scDV-AF with three AF sources—gnomAD, gnomAD+filters, and 1KGP (red shades), six-channel scDV models with and without checkpoint initialization (blue shades), and classical callers GATK HaplotypeCaller and Strelka2 (gray shades). Error bars represent standard deviation across the seven samples. All evaluations restricted to sites with ≥3× read coverage.

Compared to non-AF models, AF integration improved both precision (by 0.10) and recall (by 0.06) (Figure 2B and 2C). The largest gains occurred for heterozygous variants, where scDV-AF models achieved an absolute F1 increase of more than 0.10 (Supplementary Figure 2A), consistent with the chromosome 20 test set results. The three AF sources (1KGP, gnomAD, gnomAD+filters) showed minimal performance differences (F1 range: 0.836–0.840), indicating that scDV performance is robust to the choice of AF reference.

scDV-AF achieved performance close to the LD-based method Monopogen (F1 = 0.878, precision = 0.948, recall = 0.817), with only a 0.04 F1 deficit while using population AF databases rather than full haplotype reference panels. In contrast, bulk RNA-seq variant-calling pipelines performed markedly worse on scRNA-seq (F1 = 0.506–0.646), yielding F1 gains of 0.19–0.33 for scDV-AF and underscoring the need for specialized single-cell variant callers.

External validation establishes scDV-AF as a competitive alternative for germline variant calling from scRNA-seq. The observed F1 gains over classical callers (>0.19) and six-channel scDV (0.10), together with marked improvements in heterozygous precision (0.80 to >0.90), demonstrate that AF integration provides robust benefits across cohorts and protocols and directly mitigates key challenges posed by sparse coverage and allelic imbalance in scRNA-seq data.

### Variant calling performance across sequencing depths

We evaluated how scDV and existing methods leverage increasing read evidence by stratifying candidate SNPs across minimum coverage thresholds on the external dataset (see Methods: Evaluation across coverage cutoffs). scDV-AF variant-calling accuracy improved substantially with increasing read depth, surpassing existing methods at 10× coverage and above (Figure 3). F1 scores increased from 0.84 at 3× coverage to 0.90 at 10× and 0.917 at 20× (Figure 3A). Alternative methods either exhibited depth-dependent improvements from a lower baseline performance or demonstrated no gains with increasing coverage.

**Figure 3:**
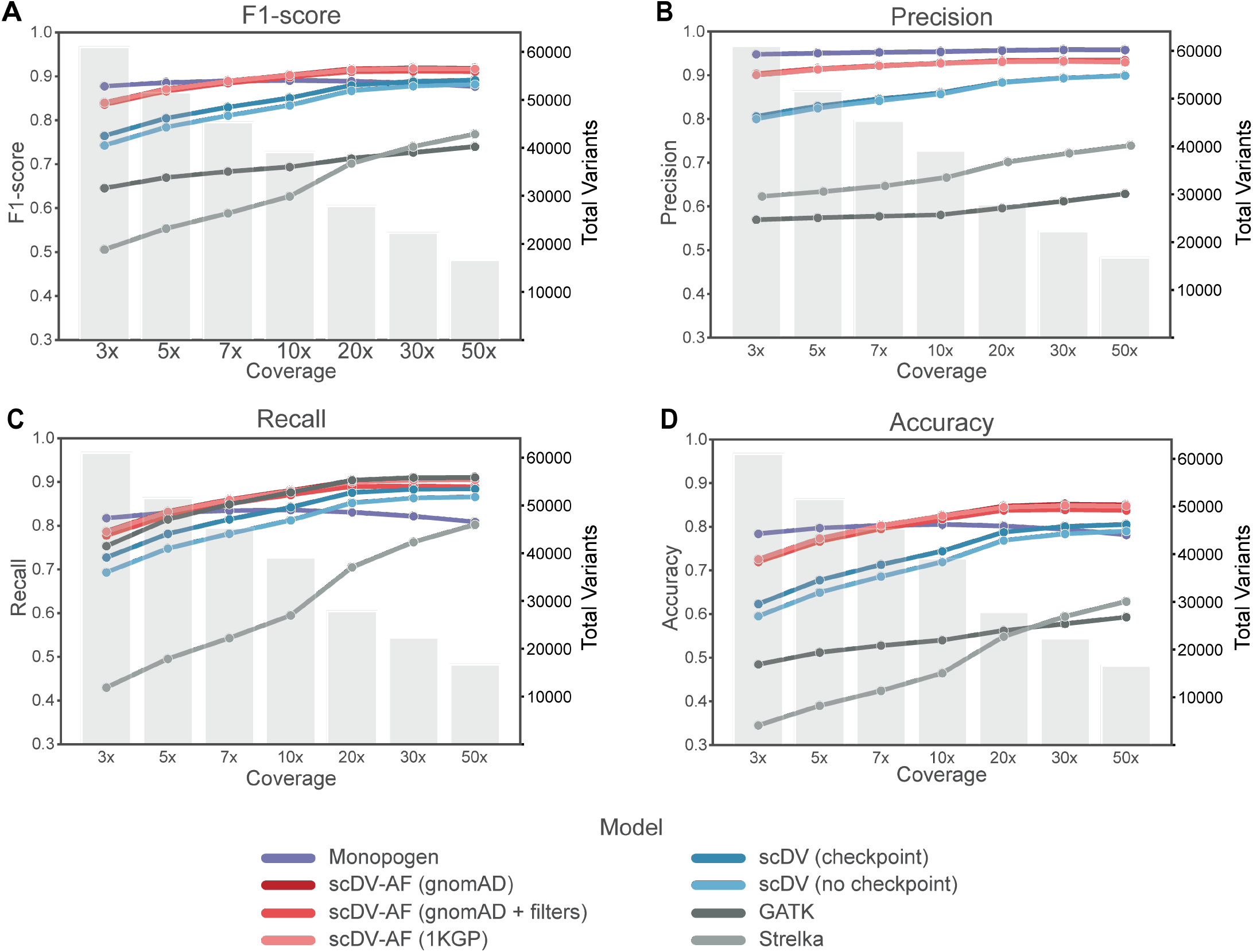
Performance metrics across minimum coverage thresholds for variant calling methods evaluated on seven independent scRNA-seq samples. Four panels show F1-score (A), precision (B), recall (C), and accuracy (D) as a function of minimum coverage cutoff (3×, 5×, 7×, 10×, 20×, 30×, 50× on x-axis). Solid lines represent six variant callers: scDV-AF models (red shades), six-channel scDV models (blue shades), Monopogen (purple), and classical callers GATK and Strelka (gray shades). The dotted gray line on the secondary y-axis shows the total number of evaluable variants at each coverage threshold. All metrics calculated only for sites meeting or exceeding the specified minimum coverage.

The improved performance of scDV-AF with increasing coverage was driven primarily by gains in recall (Figure 3C), increasing from 0.79 at 3× to 0.90 at 20×, while precision remained stable (Figure 3B). Six-channel models without AF scaled similarly with depth but from a lower baseline, indicating that AF integration provides consistent accuracy advantages across coverage levels. Monopogen performance remained stable regardless of coverage, suggesting that the current LD-based imputation approach provides limited additional benefit as read evidence increases. Classical callers improved substantially in recall but remained constrained by low precision. While the number of evaluable variants decreased with coverage thresholds (Figure 3, gray bars), approximately half of all candidate sites exceeded 10× depth, representing a substantial fraction of the data where scDV-AF achieves its strongest performance. These sensitivity gains reflect the ability of deep learning models to automatically optimize variant detection from data rather than through expert-specified statistical assumptions ^[10]^. With higher coverage, the CNN recognizes consistent alternate allele support patterns across multiple independent reads and, combined with AF priors, reliably discriminates genuine variants from technical artifacts.

Genotype-stratified analysis revealed that heterozygous variants benefited the most from increased coverage (Supplementary Figure 4). For AF models, recall improved by 0.13 from 3× to 10× while precision remained stable. Non-AF models showed even larger recall gains (0.17 at 10× reaching 0.24 at 30×) along with precision improvements (0.03 at 10× to 0.07 at 50×), suggesting that higher coverage partially compensates for the absence of population priors. Classical callers also improved substantially on heterozygous variants but remained limited by low baseline precision (reaching 0.56–0.70 at 50×). For homozygous variants (Supplementary Figure 5), all models performed well with F1 improvements up to 0.05 across coverage levels, indicating that these calls present less ambiguity regardless of AF integration.

These results demonstrate that scDV-AF leverages additional read depth more effectively than existing approaches, outperforming other methods at 10× coverage and above. The combination of data-driven pattern recognition and population-level priors enables reliable variant discrimination that scales with available read evidence, with particular benefits for heterozygous variants where allelic imbalance creates inherent ambiguity.

### Enhanced detection of rare variants in scRNA-seq data

We evaluated how scDV-AF and existing state-of-the-art methods perform in detecting rare variants, defined as variants with AF below 1% (AF < 0.01) in population databases. We compared scDV-AF against Monopogen while excluding non-AF models and classical callers because of their low performance in previous evaluations. scDV-AF outperformed Monopogen in rare variant detection, with performance advantages that progressively increased at lower AF (Figure 4).

**Figure 4:**
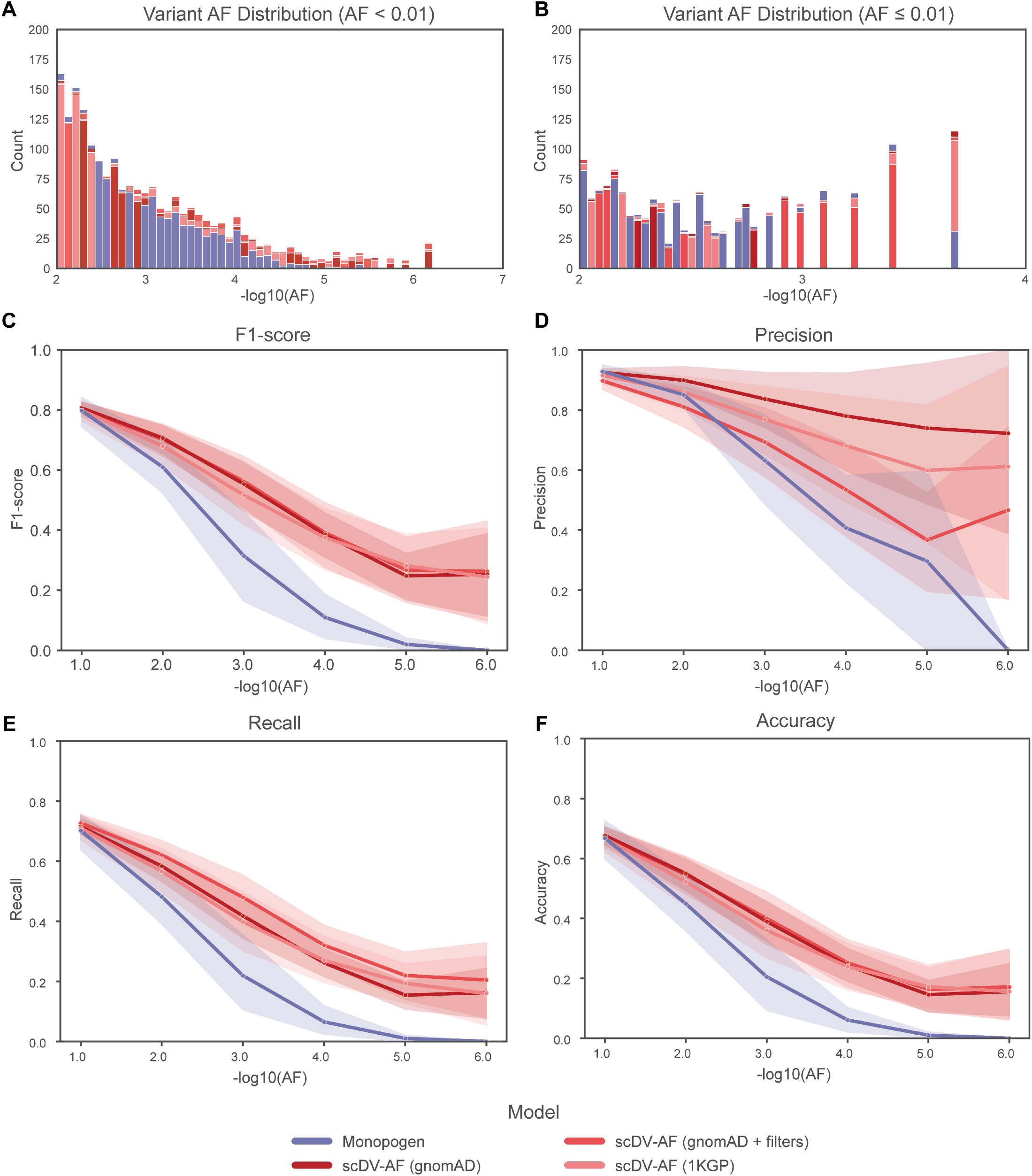
Rare variant detection in scRNA-seq. (A) Number of detected variants across gnomAD-derived AF bins in the external evaluation dataset. (B) Number of detected variants across 1KGP-derived AF bins in the same dataset. (C) F1 score, recall, and accuracy evaluated across progressively stringent AF cutoffs, highlighting performance differences for rare and ultra-rare variants. (D) Precision evaluated across the same AF cutoffs.

scDV-AF models consistently identified more true positive variants across AF bins from AF < 10^−3^ to AF < 10^−6^ in the gnomAD database (Figure 4A). At higher AF bins (AF > 10^−3^), both methods detected similar numbers of variants. However, at AF < 10^−3^, scDV-AF models consistently detected more variants, with the gap increasing progressively toward rarer alleles. Figure 4B displays analogous results using 1KGP-derived frequencies across bins from AF < 10^−2^ to AF < 10^−4^. While performance was more variable due to the smaller cohort size affecting frequency resolution, scDV-AF detected significantly more variants in the rarest bin, demonstrating robust sensitivity even with lower-resolution AF estimates.

The performance metrics confirm that higher detection rates translate into improved rare-variant detection across all measures. At AF < 10^−2^, scDV-AF gnomAD achieved F1 = 0.70 with recall = 0.62, compared to Monopogen’s F1 = 0.61 and recall = 0.48 (Figure 4C). As AF thresholds became more stringent, scDV-AF maintained relatively stable performance (F1 = 0.56 at AF < 10^−3^; F1 ≈0.39 at AF < 10^−4^), while Monopogen declined sharply (F1 = 0.31 at AF < 10^−3^; F1 = 0.11 at AF < 10^−4^). The performance divergence became even more pronounced at extremely rare AF levels: at AF < 10^−5^, scDV-AF models maintained meaningful detection capability (F1 = 0.27–0.28) while Monopogen performance collapsed (F1 = 0.02), and at AF < 10^−6^, Monopogen failed completely (F1 = 0.00), whereas scDV-AF gnomAD retained F1 = 0.26 with precision ≈0.70. The scDV-AF gnomAD model maintained precision ≈0.80 at AF < 10^−4^ and ≈0.70 even at AF < 10^−6^, whereas Monopogen precision dropped below 0.5 starting at AF < 10^−2^ (Figure 4D). When evaluating performance using 1KGP-derived AF (Supplementary Figure 6), scDV-AF models consistently outperformed Monopogen across AF thresholds < 10^−2^, although the performance gap narrowed because of less accurate AF estimates from the smaller reference population. Among AF-integrated models, scDV-AF gnomAD achieved the most stable precision-recall balance, indicating that broader population-level AF information, even from variants that did not pass all quality filters, provides better-calibrated priors than more restrictive filtering strategies. This reflects scDV-AF’s ability to evaluate each site independently using read evidence and learned AF priors, without requiring rare variants to be tagged by common haplotype markers.

These results establish rare variant detection as a key capability where scDV-AF provides substantial advantages over LD-based methods. By evaluating read evidence directly with learned population priors rather than relying on haplotype structure, scDV-AF recovered approximately 5× more ultra-rare variants (AF < 0.0001) while maintaining higher performance across all metrics at rare frequency thresholds. This finding extends DeepVariant’s established strengths in rare variant detection from bulk WES/WGS to the sparse single-cell RNA context.

## Discussion

We introduce scDV, which adapts the DeepVariant architecture for germline variant calling in scRNA-seq data and incorporates population AF information as an additional input channel, an approach not previously explored in bulk RNA-seq variant calling. While the original DeepVariant WGS/WES models incorporated AF from 1KGP, we extend this by systematically evaluating multiple AF sources, including 1KGP and gnomAD v3, using AF derived from PASS-filtered variants versus AF derived from the full gnomAD callset. These results demonstrate that population-scale priors consistently improve performance across scRNA-seq datasets. Our benchmarking establishes scDV as a strong tool for rare variant detection (AF < 0.01), where it substantially outperforms existing methods, with the performance advantage increasing progressively at lower AF. This reflects scDV’s ability to evaluate read evidence directly using learned population priors rather than relying on haplotype structure, enabling reliable detection even when rare variants are absent or underrepresented in LD reference panels. At 3× minimum coverage thresholds, scDV-AF achieves competitive performance within 0.04 F1 of Monopogen while using only AF databases rather than full haplotype panels. scDV surpasses all competing methods at 10× coverage and above, reflecting the data-driven nature of deep learning models that can effectively leverage additional read evidence.

Promising avenues for improving scDV performance include expanding training data, incorporating ancestry-specific AF, and developing hybrid approaches that integrate LD-based refinement. First, expanding to over 500 donors with paired single-cell transcriptomic and genomic variant data, including 424 donors from the full ROSMAP resource (Synapse: syn18485175) ^[24]^, 16 donors from the GTEx project ^[25]^, and additional datasets from epithelial (GEO: GSE221390) ^[26]^ and cardiac tissues ^[6]^, would improve robustness across multiple tissues and disease contexts by leveraging a substantially larger training set. Second, future work could investigate the use of ancestry-specific AF to improve variant-calling accuracy in diverse populations, as population-specific AF priors may better discriminate genuine variants from technical artifacts in samples from underrepresented ancestries. Finally, hybrid approaches that combine scDV’s direct read evaluation with LD-based genotype refinement could leverage the complementary strengths of both strategies for different variant classes. LD-based imputation methods like Monopogen may provide more accurate genotype calls for common variants when read coverage is limited ^[27]^, while scDV-AF excels at rare variant discovery where population priors and direct read evaluation are critical. Integrating both approaches could optimize genotyping accuracy across the entire AF spectrum, combining the precision of deep learning for rare variants with the power of haplotype-based inference for common variants.

## Data Availability

Model weights are available on Hugging Face (https://huggingface.co/StaticMaster/scDeepVariant), and source code is available on GitHub (https://github.com/Ilia-Buralkin/scDeepVariant).

The snRNA-seq and WGS ROSMAP dataset used for training and testing is available at https://doi.org/10.7303/syn2580853.

For external evaluation, five matched normal skin samples (P3, P5, P6, P9, and P10) with paired scRNA-seq and WES were obtained from a cutaneous squamous cell carcinoma cohort^[20]^ (scRNA-seq: GEO accession GSE144236, samples GSM4284230, GSM4284235, GSM4284237, GSM4284245, and GSM4284247; WES: GEO accession GSE144237, samples GSM4284253, GSM4284257, GSM4284259, GSM4284265, and GSM4284267).

Additional matched normal samples from esophageal squamous cell carcinoma (ESCC) patients were obtained from the ESCC single-cell study by Zhang et al.^[18]^ using two publicly deposited datasets: (i) adjacent-normal scRNA-seq data from the ESCC single-cell atlas (GEO accession GSE160269; BioProject PRJNA672851; GSA-Human HRA000195; samples GSM4870320 and GSM4870386), and (ii) matched germline WES of peripheral blood from the same patients (SRA study SRP327447; BioProject PRJNA744411; experiments SRX11378604 and SRX11378605).

## Supplementary Data

Supplementary Data are available at NAR Online.

## Acknowledgements

During the preparation of this manuscript, the authors utilized OpenAI’s ChatGPT to assist with grammar correction and language refinement. All intellectual content is solely that of the authors.

## Author Contributions Statement

Ilia Buralkin: Conceptualization (Lead); Data curation (Lead); Formal analysis (Lead); Investigation (Lead); Methodology (Lead); Software (Lead); Validation (Lead); Visualization (Lead); Writing – original draft (Lead); Writing – review & editing (Lead). Hu Chen: Conceptualization (Supporting); Data curation (Supporting); Formal analysis (Supporting); Investigation (Supporting); Supervision (Supporting); Writing – review & editing (Supporting). Junseok Park: Investigation (Equal); Project administration (Equal); Supervision (Lead); Visualization (Equal); Writing – original draft (Equal); Writing – review & editing (Equal). Zhandong Liu: Funding acquisition (Lead); Project administration (Equal); Resources (Lead); Supervision (Equal); Writing – review & editing (Supporting).

## Funding

This work was supported by the Chao Endowment and the Huffington Foundation.

## Conflict of Interest Disclosure

The authors declare no conflict of interest.

## Supplementary Figures

**Supplementary Figure 1:**
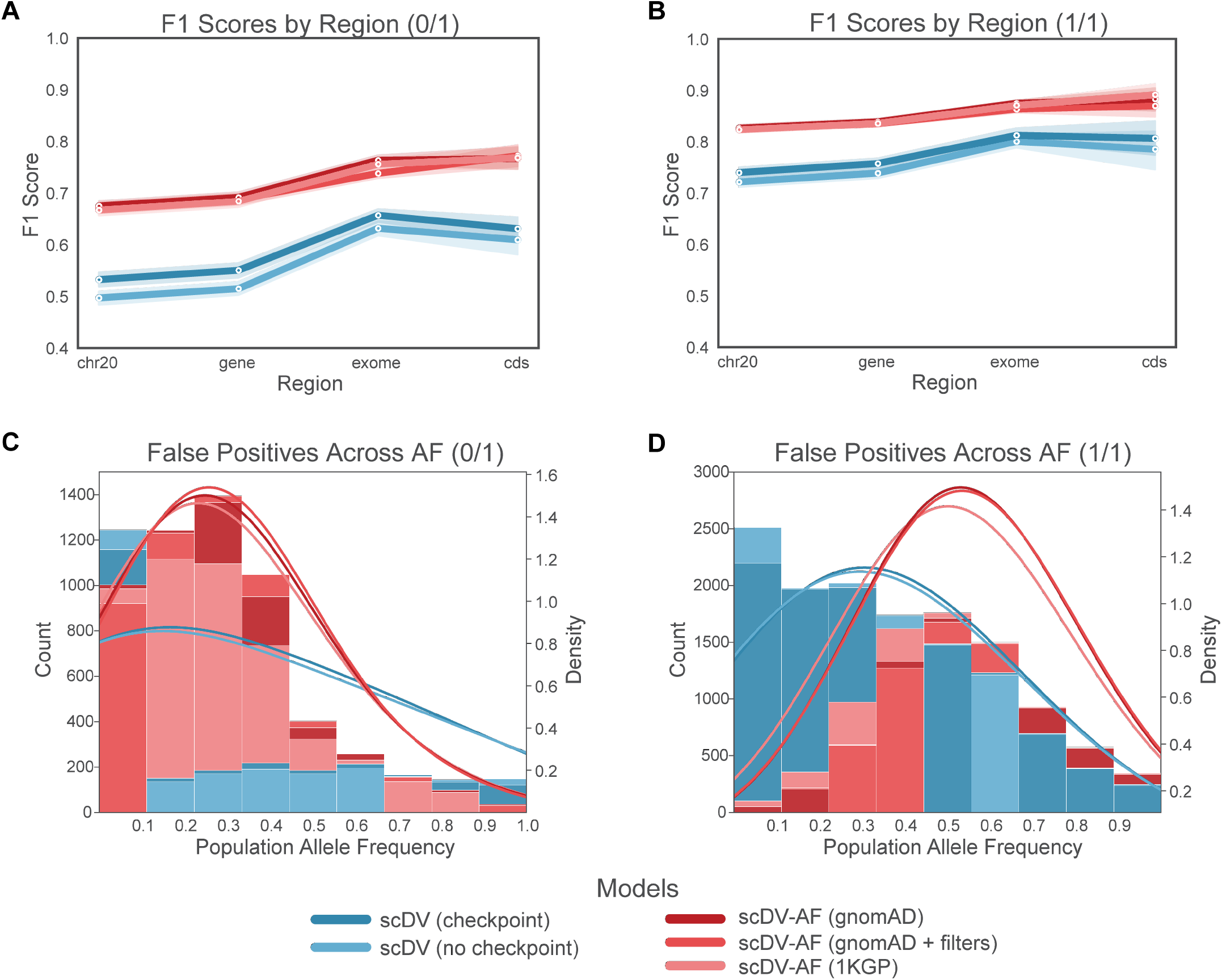
Performance of scDV models on the ROSMAP chromosome 20 holdout set, stratified by zygosity. Panels A and B show F1 scores across genomic regions for heterozygous (A) and homozygous (B) variants, respectively. Panels C and D show kernel density estimates of population allele frequencies for false positive calls in heterozygous (C) and homozygous (D) variants.

**Supplementary Figure 2:**
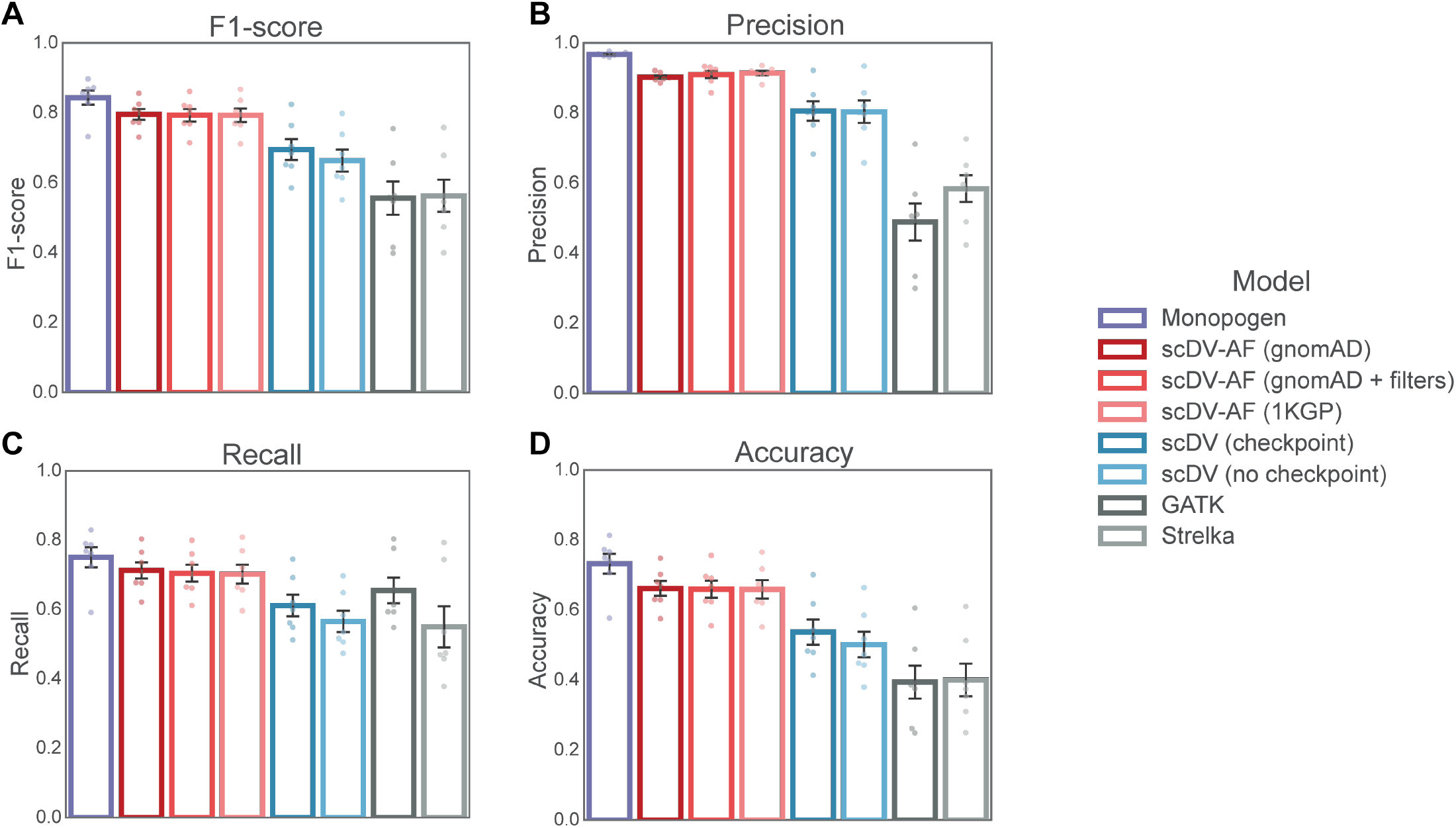
Performance on heterozygous (0/1) variants..

**Supplementary Figure 3:**
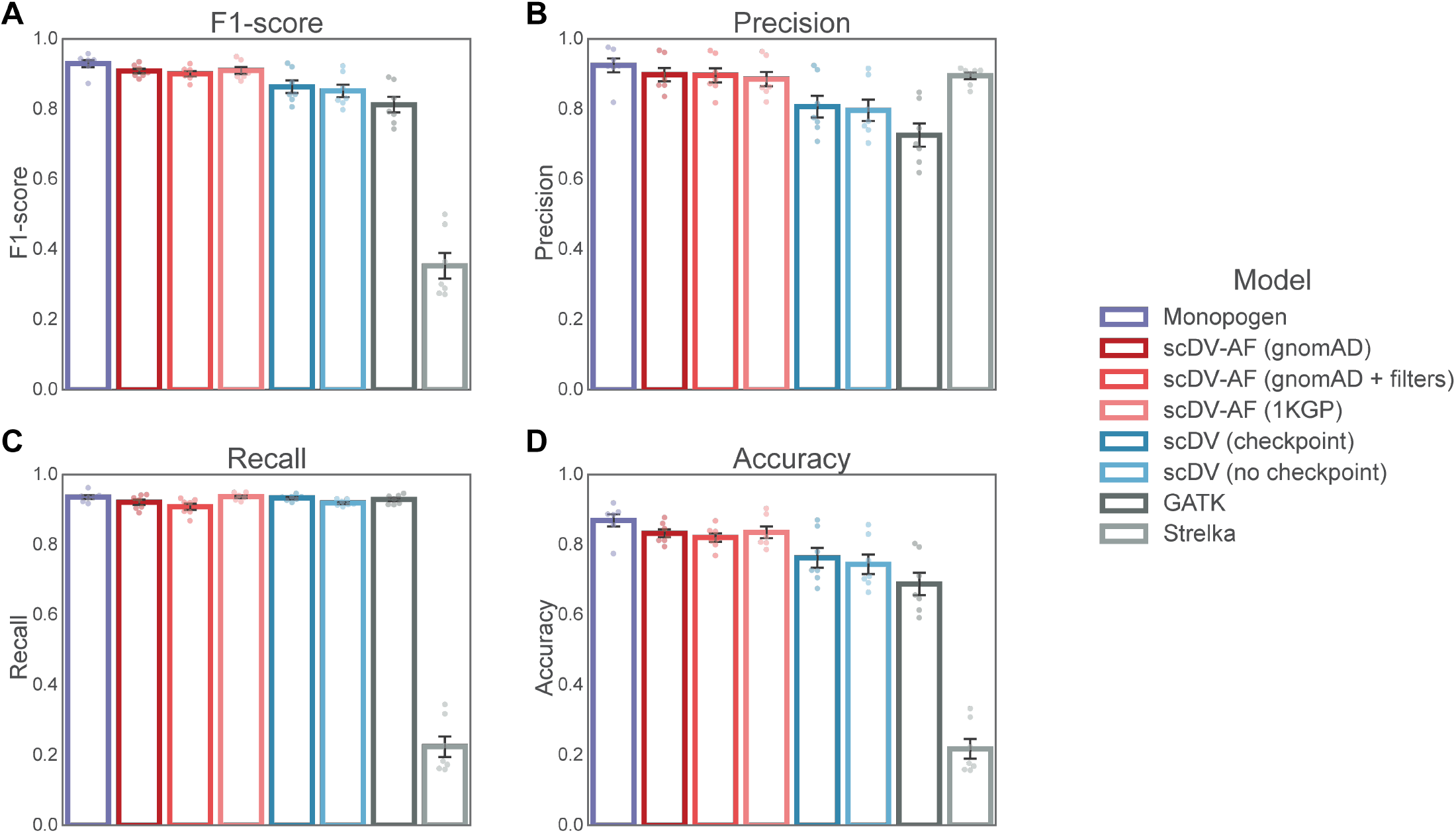
Performance on homozygous (1/1) variants.

**Supplementary Figure 4:**
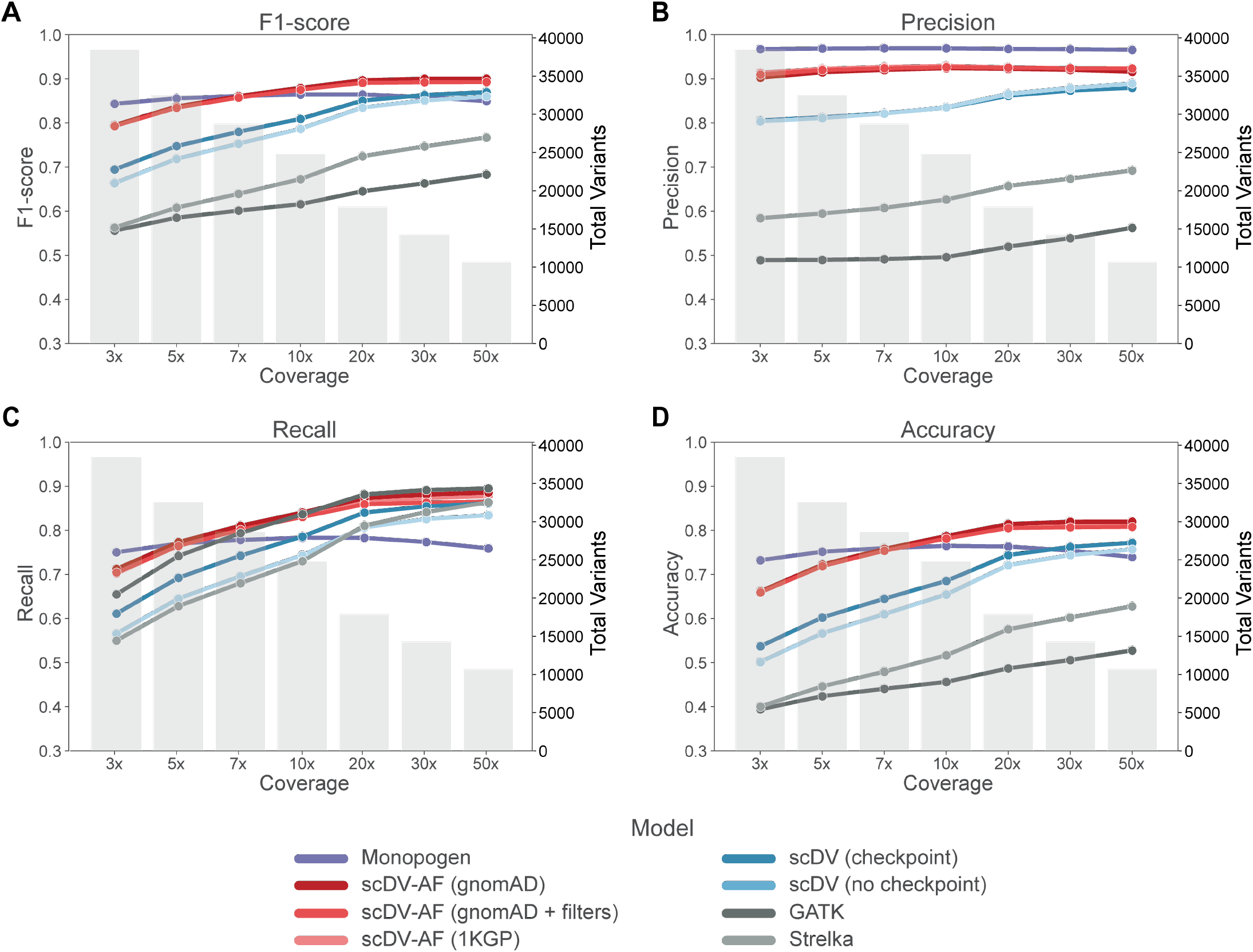
Performance metrics across coverage thresholds for heterozygous (0/1) variants.

**Supplementary Figure 5:**
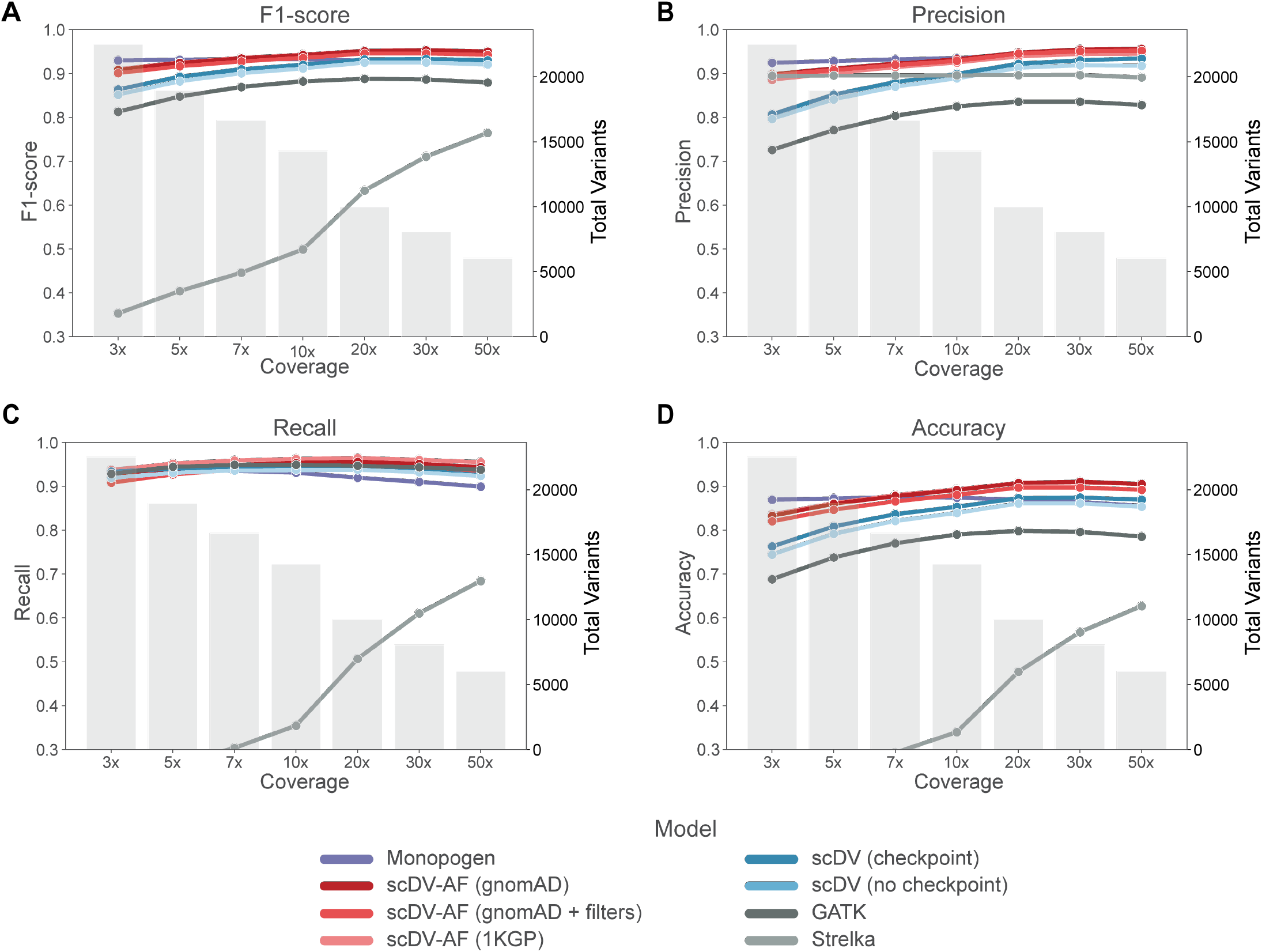
Performance metrics across coverage thresholds for homozygous (1/1) variants.

**Supplementary Figure 6:**
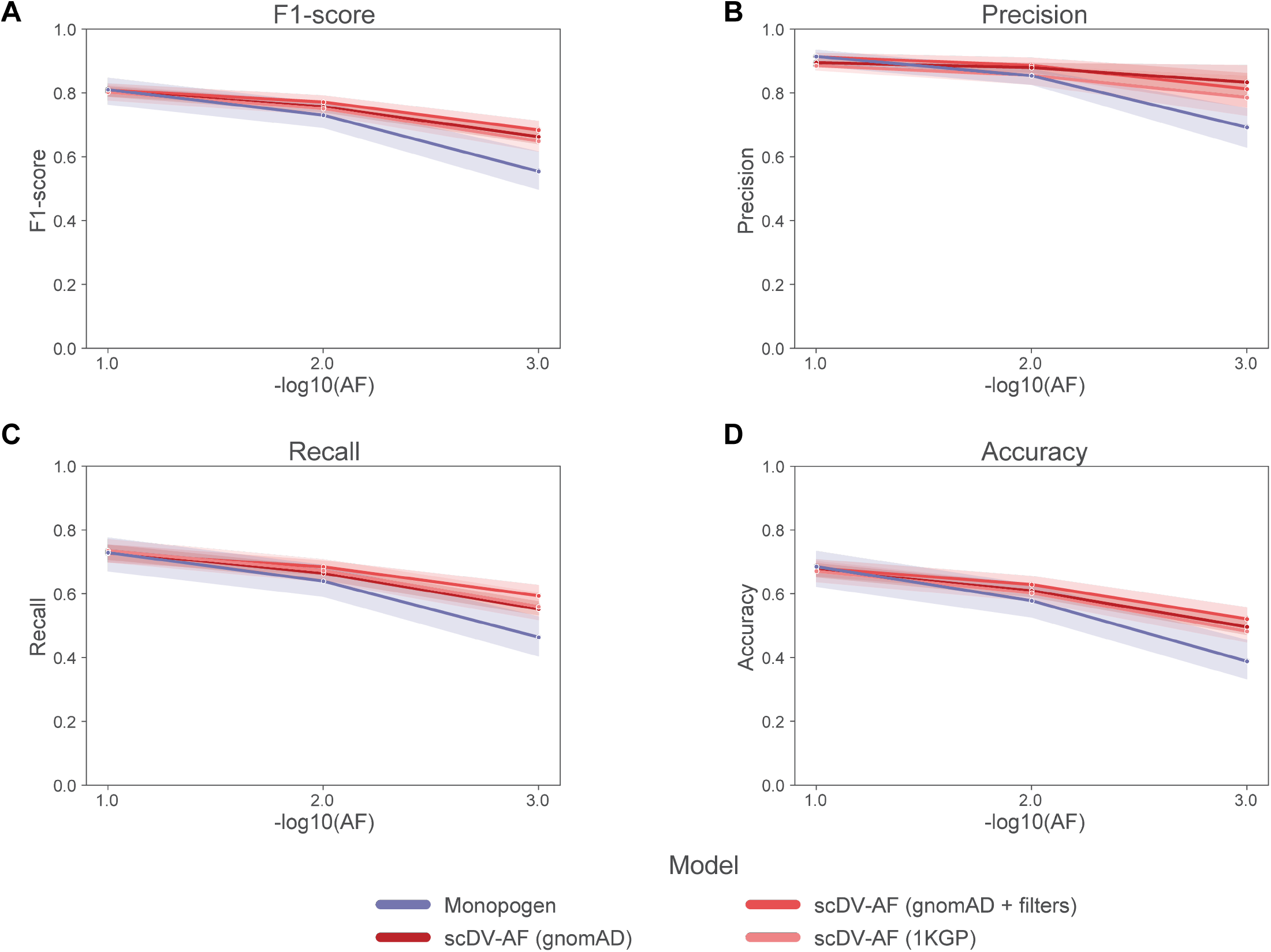
Performance metrics across 1KGP AF thresholds.

**Supplementary Figure 7:**
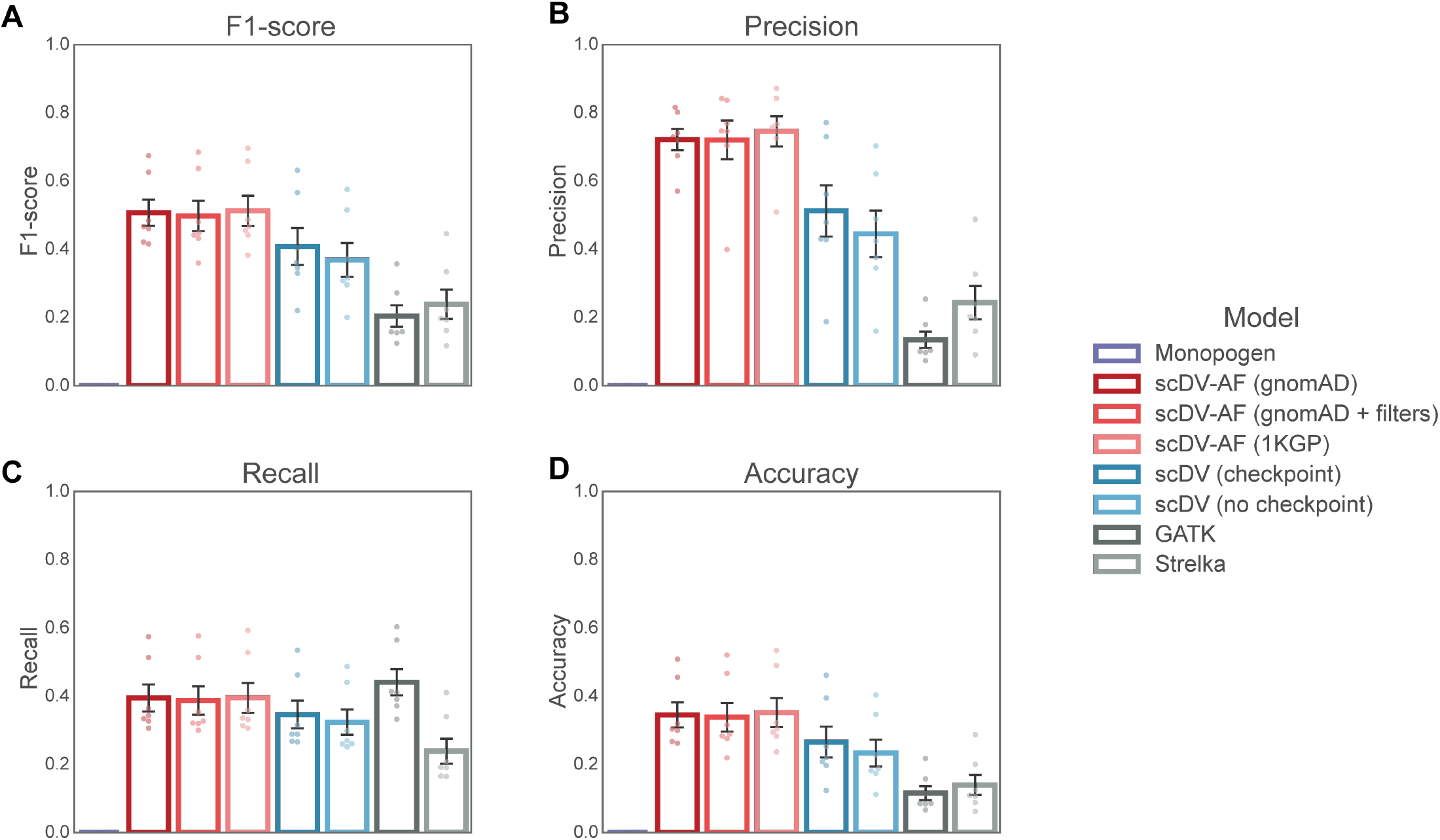
INDEL performance.

## Notes

### Competing Interest Statement

The authors have declared no competing interest.

### Summary of Updates

Author list, affiliations, and manuscript formatting updated.

